# Global inequity in scientific names and who they honor

**DOI:** 10.1101/2020.08.09.243238

**Authors:** Shane DuBay, Daniela H. Palmer Droguett, Natalia C. Piland

## Abstract

As a cornerstone of biodiversity science, Linnaean taxonomy has been used for almost 300 years to catalogue and organize our knowledge of the living world. In this system, the names of species themselves take on additional functions, such as describing features of the organism or honoring individuals. Here, we analyze the connections between bird species descriptions and who they honor from 1950 to 2019 within a context of global structures of power and access to science to interrogate how authority over the natural world is designated through Western scientific naming practices. We find that 95% of bird species described during this period occur in the Global South, but these species are disproportionately described by and named in honor of individuals from the Global North. We also find an increase through time in authors from the Global South, but Global North authors continue to disproportionately hold first author positions. Our findings show how research and labor in the Global South continue to be disproportionately translated into power and authority in the Global North, upholding and re-enacting imperialistic structures of domination. Addressing these inequities as a scientific community will require reflection and collective dialogue on the social foundations and impacts of our science.

*For working definitions of key terms, see Table 1. For a Spanish language version of the manuscript, see Supplement (para la versión en español, ver el Suplemento)*.

## 1. INTRODUCTION

The act of naming and ordering the living world cuts across cultures and language, and is an integral part of how we make sense of the world around us [1]. By naming and classifying organisms, we build a foundation for understanding their biology, which has enabled scientists to study variation and diversity [2], define biological units to conserve [3], and commodify or extirpate species [4,5]. The world as we know it is a direct result of our ability to name and catalog the natural world.

In 1753, Carl Linnaeus codified a binomial system of taxonomy, which has since become a cornerstone of Western biology and biodiversity science [6]. While the primary function of Linnaean taxonomy is to document and organize knowledge of the living world, the names of species themselves take on additional functions, such as describing features of the organism (like where it is found or what it looks like), or honoring individuals. For example, in 2017, a bird species was described from an outlying ridge of the Peruvian Andes. The bird was given the scientific name *Myrmoderus eowilsoni* in honor of the “Father of Biodiversity” — Edward O. Wilson [7]. In response, Wilson said that, “the idea of [having] a bird named after you is right up there with maybe the Nobel [Prize], because it’s such a rarity to have a true new species discovered, and I do take it as a great personal honor” [8]. As Wilson notes, descriptions of birds new to Western science have become rare events in the last 70 years (Figure 1A), and being the inspiration for a new species name is widely considered a great honor.

**Figure 1.**
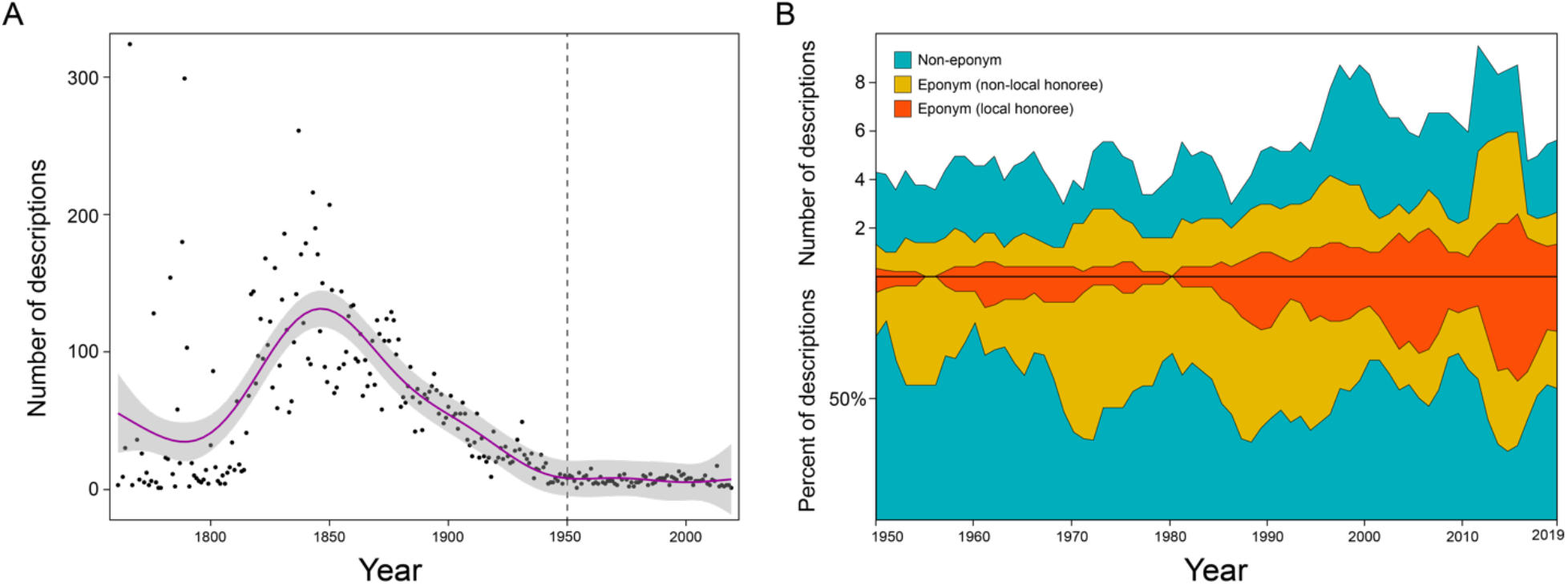
The number of bird species descriptions through time. (A) The number of descriptions by year that follow Linnaean taxonomy, starting after 1758, when Carl Linnaeus initially described 554 birds in the 10th edition of *Systema Naturae* [103]. The purple line is a LOESS regression with 95% confidence intervals (shaded gray area). The dotted vertical line marks the point in time at which the dataset for this study begins, once the magnitude of species descriptions bottoms out in the mid-twentieth century. (B) The total number (top) and percent (bottom) of descriptions split by naming category from 1950 to 2019, plotted as a five-year moving average. Species names that honor individuals (eponyms) are divided into two categories: eponyms that honor individuals from the country where the bird was described (local honoree), and eponyms that honor individuals from somewhere other than the country where the bird was described (non-local honoree). We binned all species names that are not eponyms (e.g. morphonyms, toponyms, etc.) into the third grouping (Non-eponym).

How a bird from the Peruvian Andes comes to be named after E.O. Wilson, a naturalist from the Southern U.S., can be understood through a historical lens and by considering the societal interests and global structures put in place during centuries of European and U.S. imperialism [9]. This history of European and U.S. conquest is inextricably tied to the enterprise of Western science. For example, critical advances in malaria research were funded and motivated by efforts in the late 19th century to curb European deaths in British colonies [10,11], and in 1902, Sir Ronald Ross received a Nobel Prize in Medicine for his work on the transmission of malaria, having argued that, “in the coming century, the success of imperialism will depend largely upon success with the microscope” [12]. As historian Rohan Deb Roy writes, “[Ross’] point neatly summarised how the efforts of British scientists were intertwined with their country’s attempt to conquer a quarter of the world” [11]. At present, similar dynamics are playing out in public health responses to the COVID-19 pandemic, which continue to advance imperialistic agendas by prioritizing hegemonic communities’ political power and economic well-being [13–16].

The reliance of Western science on imperialist ventures (and vice versa) is probably best catalogued in the links between naturalists and slave trade, prospecting and resource extraction, and European exploration of the 18th and 19th centuries [17–19]. The impacts of these naturalists on present-day science are difficult to overstate; the voyages of naturalists like Charles Darwin on the *Beagle* and Sir Joseph Banks on the *Endeavour* were integral parts of an imperialist enterprise [20]. Imperialism granted Western scientists unprecedented access to the world, which they translated into scientific authority, power, and wealth, fueling narratives of white supremacy, while simultaneously disregarding (or appropriating) Indigenous knowledge.

These narratives were used in turn to justify genocide, coercive labor, and exploitation, enabling the flow of material wealth and intellectual resources from colonized lands to metropoles [19,21,22]. In settler colonial states, like the U.S., nation-building has relied on European colonialist and imperialist infrastructure for global access (e.g. [23]). As a result, Western science is largely conducted through the same institutions and practices that were established to advance imperialist interests.

In this study, we interrogate the connections between imperialism and science through foundational practices like Linnaean taxonomy. Across taxonomic groups, there have been efforts to examine the historical, cultural, and ethnolinguistic roots of species names to understand how social relations shape our knowledge of biodiversity [24–38]. Here, we compiled and analyzed a global dataset of bird species descriptions and their authors to ask: who has access and power to name species, and who is honored in species names? We focus our analysis on birds because of their broad scientific and cultural relevance. We also limit our analysis to after 1950, when the opportunity to name new bird species becomes rare (Figure 1A), further intensifying the potential for recognition of authors and honorees with new descriptions. We find that global patterns of naming and authorship, extending into the present, are consistent with the historical exploitation of intellectual and material goods in the Global South [21], and advance scientific authority in the Global North, as expertise about the natural world continues to be disproportionately claimed by the Global North through publication practices. Our findings serve as a case study that reflects the inequitable structures at the core of Western biodiversity science and their resulting disparities, e.g. in access, labor, collaboration, power, and designations of expertise and authority. This study shows how historical inequity continues to shape present day research practices, and highlights the need for community-level reflection and dialogue beyond the hegemonic center.

## 2. METHODS

We compiled a dataset of bird species described within Linnaean taxonomy from 1950 to 2019. We recorded information about: the country from where each bird was described, the etymology of each species name, the authors of each description, and the journal and language in which each description was published. All data compiled for this study are from publicly available sources. With this dataset, we then assessed geographic patterns of naming and authorship across this time period.

### (a) Dataset

As our base dataset, we used the Birdlife International global avian checklist, which is regarded as a dominant authority in avian taxonomy (HBW and BirdLife Taxonomic Checklist v4 Dec 2019). Importantly, this checklist includes information on the authors and year of each species’ description. For entries from 1950 to 2019, we removed taxa that are currently recognized as species but were originally described as subspecies. We identified these entries by looking at the original species/subspecies description for each entry. Removing these taxa ensures that the level of honor at which a taxon is described is consistent between entries. Additionally, we added in taxa for which the opposite scenario is true—that is, we included taxa that were originally described as species following Linnaean taxonomy but are not currently recognized as species by the BirdLife International checklist committee. We included these “reclassified” species in the dataset because they were originally described with the intention of being at the species level. We identified these taxa using *Bird Species New to Science: Fifty Years of Avian Discoveries* [39], which reports a comprehensive list of taxa described as species between 1960 and 2015. For taxa described as species between 2015 and 2019, we conducted an internet search to identify recently described species to include in the dataset. This search was guided by the “List of bird species described in the 2010s” on Wikipedia (https://en.wikipedia.org/wiki/List_of_bird_species_described_in_the_2010s), and our knowledge of birds described during these years as members of this scientific community. Lastly, we did not include descriptions in which the species was extinct at the time of description. Figure 1A includes all (and only) entries from the HBW and BirdLife Taxonomic Checklist v4, but for all analyses and subsequent figures, subspecies and extinct taxa were removed, and “reclassified” species were included.

### (b) Data associated with species names

We defined a species’ type locality as the country from where the species was described, which we determined from locality data associated with holotype specimens. We then classified countries and island regions as either Global North or Global South—here and for author metrics and journals below—based on the United Nations classifications of “developed” and “developing” regions (https://unstats.un.org/unsd/methodology/m49/), with “developed” corresponding to the Global North, and “developing” corresponding to the Global South [40]. We use the terms Global North and Global South in the sense of Dados & Connell (2012) to place boundaries in the dataset and analyze geopolitical relationships of power and dominance [41]. We recognize, however, that this binary does not represent cultural, material, and lived experiences around the world, and that other terminologies are more appropriate to demarcate the world and its peoples for different purposes (e.g. ‘majority world’ [42]).

We classified species names based on their meaning and derivation, placing each species into one of nine naming categories defined in the *Helm Dictionary of Scientific Bird Names* [43]. The categories include: eponym—named after a person or persons; morphonym—named after morphological characteristics, like plumage; toponym—named in reference to a geographic place; autochthonym—named in an indigenous language; taxonym—named in relation to other taxa; bionym—named after habitat or environmental conditions; ergonym—named after behavioral characteristics, like breeding or display behaviors; phagonym—named after diet or prey type; and phononym—named after vocal characteristics.

We further divided eponyms into five categories: local—named after an individual from the country of the species’ type locality; non-local—named after an individual from a country other than the species’ type locality; fictional—named after a fictional character; titles—named after an honorific title, like Prince; and group—named after two or more people. To determine if an eponym was local or non-local, we had to first infer where an honoree was from, which we defined as the country where they were born, and determined using *The Eponym Dictionary of Birds* [44]. For example, the entry for Maria Koepcke says, “born Maria Emilia Ana von Milkulicz-Radecki in Leipzig, Germany,” and the entry for Alfonso Maria Olalla says, “an Ecuadorian professional collector, who lived in Brazil (mid-1930s) and took Brazilian citizenship.” We recorded the countries where they were from as Germany and Ecuador, respectively. For individuals who lack this information in *The Eponym Dictionary of Birds*, we determined where they were born from other publicly available sources, such as curriculum vitae available online or obituaries published in society journals.

We then assessed gender designations for individuals honored in eponyms, based on the Latin endings of species names (-ae = woman, -i = man, -orum = group of women/men or group of all men, -arum = group of women). It is worth noting that the Latin language and the codes that govern Linnaean taxonomy impose binary gender designations for eponymous names (ICZN Article 31.1 [animals], ICNafp Article 60.8 [plants], ICNB Appendix 9 [bacteria]), and while these rules allow us to assess the binary gender designations of eponyms, they also erase the spectrum of gender identities.

### (c) Author metrics: institutional affiliation, country of origin, but not gender

We compiled author data from the publication of each species description. We recorded the number of authors on each publication and each author’s institutional affiliation. For authors with more than one affiliation listed, we used their first institution listed for our analyses, as this institution is given and perceived as having priority. When an author’s institutional affiliation was not included in a species description, which is the case for some publications earlier in the dataset, we inferred their institutional affiliation at the time of publication (when possible) from other publicly available sources, such as obituaries.

We inferred an author’s country of origin from a combination of where they were born and where they received an undergraduate education, which we compiled from publicly available sources, such as obituaries, personal websites, curriculum vitae, etc. Our country of origin metric is intended to capture two things: the place where an individual received their formative education, and the academic conventions under which they were trained. When possible, we defined an author’s country of origin as the country where they were born (61% of authors), but if this information was unavailable, we used the country of their undergraduate institution when available (14% of authors). This combined approach helps us increase coverage, and when both data types were available for an author (34% of authors), the two metrics show 93% congruence. We then classified each author as local (the author’s country of origin is the same as the country of the species’ type locality), or non-local (the author’s country of origin is different from the country of the species’ type locality).

We deliberately refrained from inferring authors’ gender because we did not ask authors to self-identify their gender(s) (for more information on operationalizing gender as a research variable, see [45,46]).

### (d) Journal location and the language of species descriptions

For each journal in our dataset where a species description was published, we determined the home country of that journal based on the journal’s self-reported location. We also determined the language of each species description based on the language used in the title of the description.

### (e) Statistical analysis

We assessed changes through time in authorship and eponym patterns using simple linear regression, with year as a single predictor variable. We also used logistic regression to assess whether or not species descriptions have local first authors or at least one local author, and we used Poisson regression to assess changes in the number of authors on a description (both of these analyses incorporated year as a single predictor variable). We then analyzed if eponym patterns (local honoree vs. non-local honoree) were significantly predicted by the authors’ country of origin (local vs. non-local) using logistic regressions, with year as a fixed effect. All analyses were conducted in R v3.6.2 [47].

## 3. RESULTS

From 1950 to 2019, 95% of bird species have been described from countries in the Global South (n = 366 of 385). During this period, only 17 species were described from the Global North. One species had an undetermined geographic placement, as a type locality was not given with its description. Species were described from 68 countries (Figure 2). However, half of all species were described from five Global South countries alone: Brazil (n = 68), Peru (n = 57), Colombia (n = 25), Philippines (n = 23), and Indonesia (n = 20).

**Figure 2.**
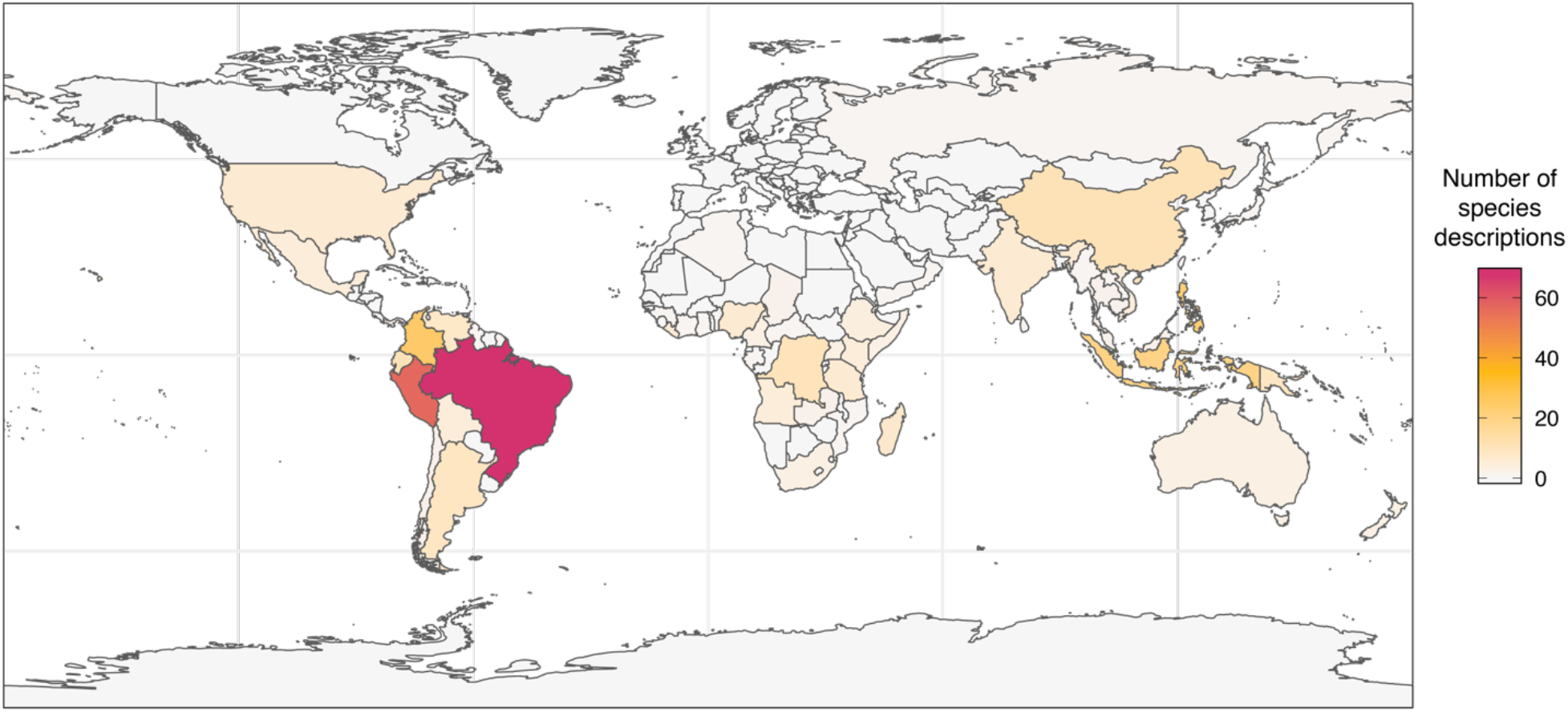
The number of bird species described from a given country from 1950 to 2019.

Half of all species described since 1950 were eponyms (50%, n = 193), i.e. named after people, like *Myrmoderus eowilsoni*. The other half of species were named after defining characteristics (50%, n = 192), like morphological features, behavior, or where it is found, like *Pyrrhura peruviana*. Over time the number of eponyms increases (R2 = 0.141, p = 0.001) as the other naming categories remain steady (R2 = 0.011, p = 0.379; Figure 1B).

### (a) Who is honored in species names?

#### (i) Eponyms disproportionately honor individuals from the Global North

Of the eponyms that are named after a single individual (n = 185), the type locality for 96% of these species are in the Global South, but the majority of these eponyms are named in honor of individuals from the Global North: 68% (n = 124) of eponyms honor individuals from the Global North, while 30% (n = 54) honor individuals from the Global South. The honoree’s country of origin was unknown for 5 eponyms. When we assessed the countries from where each bird was described, only 31% (n = 56) of eponyms honor individuals from the species’ type locality, while 67% (n = 122) of eponyms honor individuals from a different country. Additional eponyms are named after fictional characters, honorific titles, or have unknown etymology (n=7), or are named after groups of people (n = 8), of which three are named after Indigenous groups from the region where the species occurs. Although the majority of eponyms honor individuals from the Global North, we found an increase toward the present in eponyms that honor local individuals—i.e. from the Global South (R^2^ = 0.242, p< 0.001; Figure 1B).

#### (ii) Gender disparity in eponyms

Gendered patterns of naming and adherence to a binary system of gender classification capture another axis along which the imperialist/colonialist foundations of Western science manifest through research practices (for critical analyses of the links between gender and imperialism see [48–50]. Of the eponyms that honor a single individual (n =183), 81% (n = 149) honor individuals categorized as men, and 19% (n = 34) honor individuals categorized as women. The observed gender disparity in eponyms is also paired with disparities in how authors characterize honorees within the text of species descriptions. For example, honorees categorized as men are often described as *colleagues* and *friends, notable scientists*, and *patrons*, while half of all eponyms that honor individuals categorized as women describe these individuals as *wives* (n = 13) and *daughters* (n = 4). To put these differences into perspective, only one honoree categorized as a man is characterized as a son, and not a single honoree categorized as a man is characterized as a husband. This disparity is further reflected in the fact that 59% (n = 20) of eponyms that honor individuals categorized as women use only their given name, while 12% (n = 4) use given name and surname, and 29% (n = 10) use only surname. In contrast, only 1% (n = 2) of eponyms that honor individuals categorized as men use only their given name, while 2% (n = 3) use given name and surname, and 97% (n = 144) use only surname.

### (b) Who has access and power to name species?

#### (i) Authors are disproportionately from the Global North

From 1950 to 2019, 545 individuals authored the 385 bird species descriptions, filling a total of 1012 author positions (descriptions have anywhere from 1 to 16 authors). We were able to compile country of origin data for 76% of all authors (n = 412 of 545) and for 84% of total author positions (n = 848 of 1012). Of these authors, 62% (n = 255) are from the Global North and 38% (n = 157) are from the Global South. We also recorded the institutional affiliation (at the time of the species description) for 98% of total author positions, which shows a similar pattern: 60% of authors were affiliated with institutions in the Global North (n = 595), and 40% were affiliated with institutions in the Global South (n = 393). These results show that the majority of authors are from the Global North and affiliated with Global North institutions.

To assess the potential impacts of having 24% (n = 133 of 545) of authors with missing country of origin data, we examined the connections between an author’s birthplace and their institutional affiliation(s). For individuals in the dataset with known birthplace and institutional affiliation(s) (which accounts for 59% of authors), 72% (n = 236) of these authors were born in the same country as the institutions where they worked, and 7% (n = 23) of authors were born in the same country as at least one of the institutions where they worked (i.e. these 23 authors were affiliated with multiple institutions in different countries). Of the remaining 21% of authors (n = 67) who were born in a country that was different from the institutions where they worked, this movement was largely within the Global North (33%, n = 22) or from the Global North to Global South institutions (49%, n = 33). We documented one instance of movement within the Global South, and only 16% of authors (n = 11) who shifted countries moved from the Global South to Global North institutions. Given the much higher prevalence of movement within the Global North and from the Global North to Global South institutions, these data suggest that the percent of authors whose country of origin is in the Global North should be higher than the percent of authors whose institutional affiliation (for which our dataset is 98% complete) is in the Global North. Thus, the country of origin data analyzed in this study likely underestimate the number of authors from the Global North.

#### (ii) First authors, author lists, and authority

In the biological sciences, the first author on a publication typically receives the most credit for the work, and is viewed as a primary authority on a publication’s contents [51,52]. We therefore examined metrics for first authors to explore who is perceived by the scientific community as the authority on a species description. We found that 71% of first authors (n = 275) are from the Global North, and 21% (n = 82) are from the Global South. The country of origin for 7% of first authors (n = 28) was unknown. In 55% of the cases where the first author is from the Global South (n = 45), all authors on the description are from the Global South. The first author’s country of origin was different from the species’ type locality for 72% of species descriptions (n = 276), and the same for 22% of species descriptions (n = 85). The first author’s country of origin was unknown for the remaining 6% of descriptions (n = 24) (for one description the species’ type locality was unknown). The prevalence of first authors from the Global South increases significantly toward the present (R^2^ = 0.143, p = 0.014), but the prevalence of first authors from the Global North remains consistent (R^2^ = 6.749e-5, p = 0.947) and is always higher (Figure 3A).

**Figure 3.**
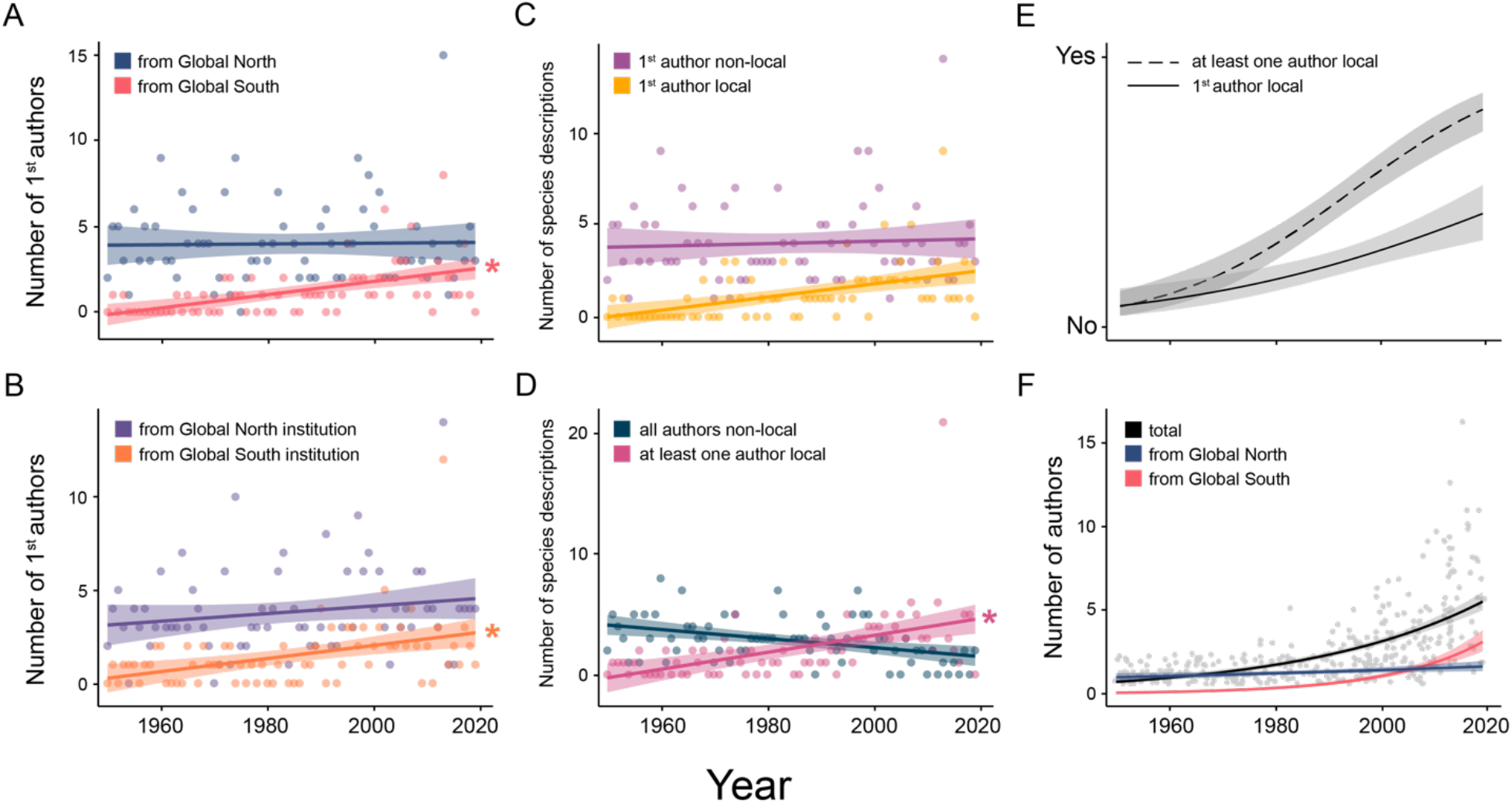
Authors’ country of origin and institutional affiliation through time. (A) The number of first authors for a given year whose country of origin is in the Global North vs. Global South. (B) The number of first authors for a given year whose institutional affiliation is in the Global North vs. Global South. (C) The number of species descriptions for a given year in which the first author’s country of origin is the same as the species’ type locality (1st author local) or different (1st author non-local). (D) The number of species descriptions for a given year in which at least one author is from the species’ type locality (at least one author local) vs. when not a single author is from the species’ type locality (all authors non-local). In panels A-D, (*) denote significant changes (p < 0.05) in the response variable through time from simple linear models. Each point is the number of descriptions in a given year for each category. (E) Logistic regressions of the data from panels C (solid line) and D (dotted line), showing changes through time in whether or not species descriptions have local first authors or at least one local author. (F) The total number of authors on a description (total), and the number of authors on a description in which an authors’ country of origin is in the Global North or Global South. Lines plot Poisson regressions. Each gray point is the number of authors on each species description, and for Global North and Global South regressions we only used descriptions for which we had country of origin data for all authors, which represents 81% of total descriptions. For all panels, shaded areas are 95% confidence intervals for each regression.

When we looked at institutional affiliation, 70% of first authors (n = 268) were affiliated with institutions in the Global North, 27% of first authors (n = 104) were affiliated with institutions in the Global South, and the institutional affiliation was unknown for 3% of first authors (n = 13). Similar to the authors’ country of origin patterns, the prevalence of first authors from Global South institutions increases significantly toward the present (R^2^ = 0.142, p = 0.007), but the prevalence of first authors from Global North institutions remains consistent (R^2^ = 0.018, p = 0.282) and is always higher (Figure 3B). Taken together, these data show that the perceived authorities on species described from the Global South are largely scientists from the Global North.

We found an increase (though non-significant) through time in the percent of species descriptions for which the first author’s country of origin is the same as the country of the species’ type locality (R^2^ = 0.053, p = 0.137; Figure 3C). However, for most descriptions, the first author’s country of origin was different from the species’ type locality (Figure 3C,E). In fact, for 51% of descriptions (n = 195), not a single author was from the species’ type locality. For 39% (n = 149) of descriptions at least one author was from the species’ type locality, and this information was unknown for the remaining 11% of descriptions (n = 41). Toward the present, however, we found a significant increase in descriptions that include at least one author from the species’ type locality (R^2^ = 0.189, p = 0.002), and a near-significant decrease in descriptions without a single author from the species’ type locality (R^2^ = 0.056, p = 0.068; Figure 3D), resulting in a pattern inversion in the 1990s (Figure 3E). That is, before 1990, most author lists were exclusively non-local, while after 1990, most author lists included at least one author from the species’ type locality (Figure 3D,E).

When we look at the entire author list for a species description, the number of authors increases significantly toward the present (GLM: χ^2^ = 288.60, p < 0.001; Figure 3F), which appears to be driven by the addition of authors from the Global South (Figure 3F), rather than changes in the number of authors from the Global North, which remains relatively stable through time (Figure 3F). This result reflects a significant increase toward the present in authors from the Global South (GLM: χ^2^ = 200.28, p < 0.001; Figure 3F).

### (c) Impacts of an author’s country of origin on eponym patterns

When the first author’s country of origin is consistent with the species’ type locality, the species is 62% more likely to be named in honor of someone from that country (GLM: χ^2^ = 18.68, p < 0.001; Figure 4). Regardless of first authorship, however, if there is at least one author whose country of origin matches the species’ type locality, then the species is 47% more likely to be named in honor of someone from that country (GLM: χ^2^ = 21.88, p < 0.001; Figure 4).

**Figure 4.**
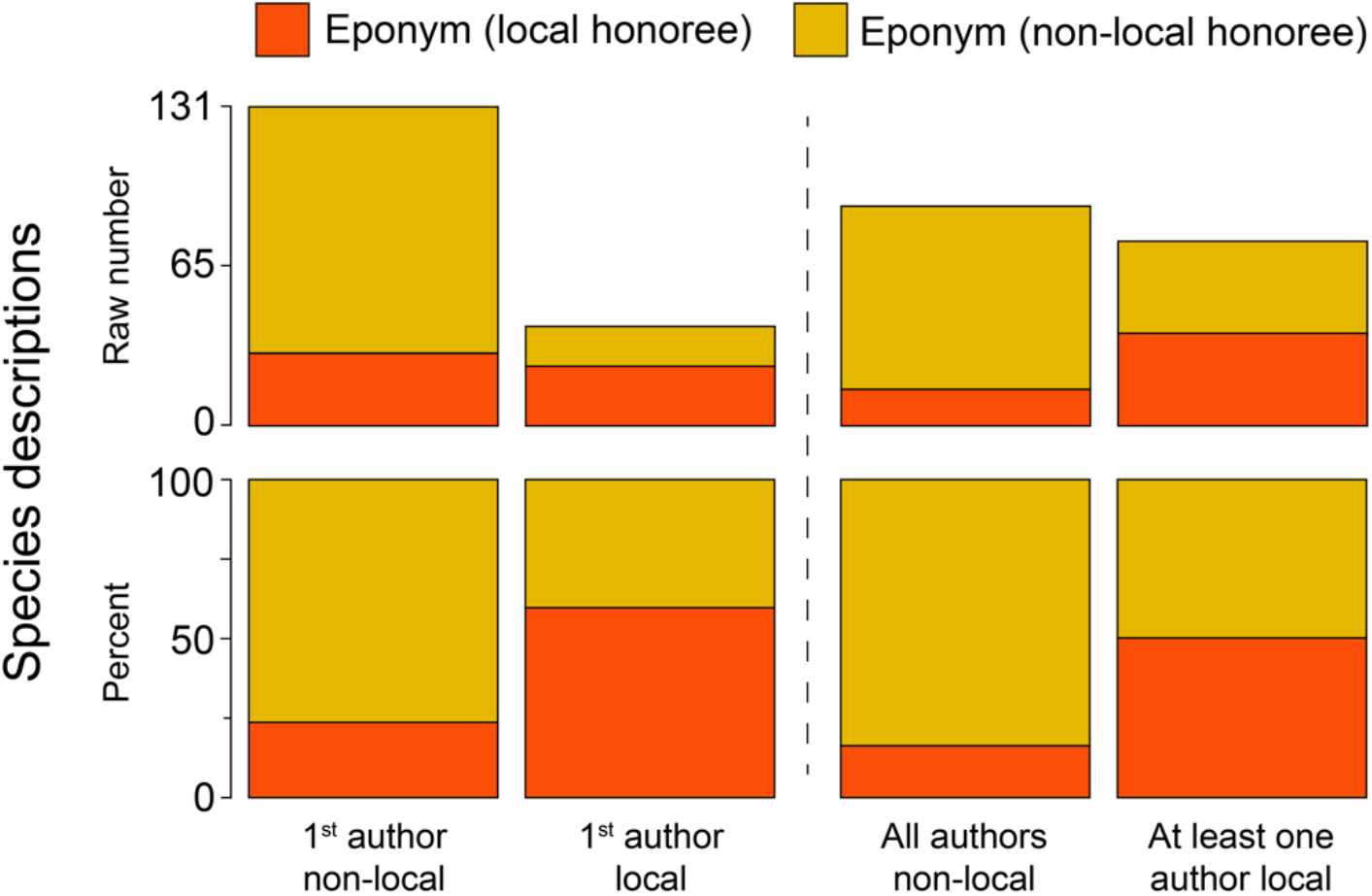
Eponym patterns based on an author’s country of origin, comparing eponyms that honor individuals from the country where the bird was described (local honoree), and eponyms that honor individuals from somewhere other than the country where the bird was described (non-local honoree). The four author classifications follow the classifications in Figure 3B,E. The top plots show the raw number of species descriptions in each category. The bottom plots show the percent of species descriptions in that category.

### (d) Journals and the language of species descriptions

We found that 85% of species descriptions are published in journals based in the Global North (n = 329), 13% of descriptions are published in journals based in the Global South (n = 51), and 1% of descriptions are published in journals with unknown geographic placement (n = 5). We found that 70% of descriptions are published in journals that are based in countries where English is recognized as an official language (n = 268), but 82% of descriptions are written in English (n = 316). This excess of English-language descriptions consists of 57 descriptions written in English that are published in journals based in countries where English is not an official language. The other languages of species descriptions are: Portuguese (n = 20), French (n = 18), German (n = 14), Spanish (n = 9), Russian (n = 1), and Vietnamese (n = 1). We were unable to classify language for six species descriptions because we lacked access to these publications.

## 4. DISCUSSION

Our results show how a foundational practice in Western science still adheres to global structures of access and power that disproportionately benefit the Global North. As professional scientists from the Global North who are affiliated with Global North institutions, we (the authors) have had access to funding and career opportunities in science that are the product of the wealth amassed through genocide, coercive labor, land seizure, and resource extraction by the U.S. and Britain (e.g. see [53]; please see our statement of Land Acknowledgement below).

Our positionalities and perspectives as Global North researchers inform our decision to focus our critique below on the hegemonic center that we are a part of. In this critique we refrain from defining solutions or best practices, and instead recommend self-reflection on positionality and power, and direct dialogue with communities beyond the hegemonic center.

### (a) Inclusion, access, and power within Western science

The patterns of authorship we observed show that researchers from the Global South have increasing opportunities to participate in Western science (Figure 3), which appears to impact naming outcomes (see Figure 4). We see an increase through time in the number of authors on a description, which is driven almost exclusively by an increase in authors from the Global South (Figure 3F). This formal inclusion of Global South authors, however, does not broadly translate into Western metrics of primary authority, like first authorship (compare the differences between Figure 3C and D, and between Figure 3A and F). As a result, Western scientific authority continues to be consolidated in the Global North. The increase in Global South authors tracks the efforts in recent years by Global North institutions to expand participation in Western science [54], and also tracks the recent increase in international collaborations [55]. These two academic trends have been motivated by a model in which diversity and inclusion equal better science, higher rankings, and increased marketability [56–58]. These initiatives, however, are documented to be largely symbolic, utilizing labor and collaborations to serve academic markets in the Global North and legitimize authority and dominance structures already in place [59–65]. This model of inclusion and collaboration prioritizes the Global North’s power to theorize and conceptualize (e.g. the scientist that extrapolates the observation of an individual bird to the *naming of a full species)*, while relying on the Global South to connect the work to the material world (e.g. the local guide/resident/scientist who facilitates the physical work to *find the individual bird*) [66]. Furthermore, this model of inclusion frames the value of people and their perspectives in terms of how they can benefit those currently in power, without challenging those power structures (e.g. [67]).

The continued dominance of English as the *lingua franca* of academic science is another way that makes concrete the power imbalance between the Global North and Global South [68]. Working within a highly commercialized system of science publishing and communication, the English learner experiences a disproportionate burden, as they must learn this additional language to publish in high impact and international journals (i.e. journals with higher impact factors) to advance their careers [69–71]. This expectation of English is highlighted in our dataset, for example, in the higher percentage of descriptions written in English (82%) than the percentage of descriptions published in journals based in English-speaking countries (70%). Getting to the point where an English learner can write a species description assumes not only that the researcher has something to write about (e.g. a bird), but that the researcher has had access to English classes and/or English-speaking colleagues/contacts/editing services, by way of financial means, time, a global social network, or institutional support, thus exacerbating global and within-country socioeconomic and power inequities [71]. For academic science that challenges, rather than perpetuates, global power inequities, we must reassess current pressures for a monolinguistic system [70–73].

It is essential to acknowledge that this study looks at knowledge production, access, and power within Western naming practices *from the perspective of the Global North*. Implicit in this perspective is that first authorship, and authorship of publications in general, are ways to establish authority and accumulate power in knowledge production, which in itself is worth questioning. For example, how do established authorship norms promote inequity and dominance in Western science (e.g. [74])? Furthermore, while our work examines dynamics between the Global North and South within a Western context, these dynamics are mirrored in broader structures of dominance between Western science and Indigenous science (as defined by [75]). The imperialistic dynamics that created the current structures of access and power within Western science between the Global North and South have also enabled Western science to assert dominance in global knowledge production [76], while erasing, appropriating, and subjugating Indigenous knowledge and authority. Solutions to build a more equitable global scientific community—if that is in fact our goal within Western institutions—will ultimately require actions that redress current structures of dominance and authority built on dispossession, violence, and white supremacy.

### (b) Naming and authorship across clades and the perceived value of taxonomy

In this study, we focus on a specific subset of scientific publications, taxonomic descriptions of birds, which raises questions about patterns of authorship and naming in other clades, and the position of taxonomic descriptions (and their perceived value) within the biodiversity literature. We focus on bird descriptions because of their relative rarity and the broad importance of bird research in the scientific literature. It would be informative to know if the patterns we observe for bird descriptions are similar in other clades given that the geopolitical backdrop of taxonomic science is shared across clades. However, we might expect patterns to vary based on clade-specific qualities, like species abundance, the perceived cultural/societal value of a group of organisms, the environments in which the organisms are found (and ease of accessing those environments), and gatekeeping by researchers studying those organisms. Looking at authorship and naming patterns across clades would shed light on how clade-specific cultural and biogeographic features shape how power and authority manifest through research practices.

Although species descriptions are foundational for biodiversity science, their overall perceived value within the Western scientific community has shifted in recent decades (see [69,77–81]). This shift is consequential for understanding the trends we observe in authorship. Across the timeframe of our dataset, biodiversity science has been transformed by major technological advancements, like the advent of DNA sequencing and more recently by high-throughput sequencing methods. With these technologies, institutional research priorities and incentives have shifted, accompanied by a decline in the perceived prestige and value of taxonomic work [77,82]. Taxonomy is now widely considered a declining or dying science, not because taxonomic work is no longer relevant to biodiversity science and conservation, but because it has been devalued in the Global North as a research priority and viable profession [82,83]. Paradoxically, as taxonomic work has been devalued, biodiversity research and global research priorities continue to be grounded in, and reliant upon taxonomic labor [69,84]. What does this mean for individuals doing taxonomic work?

In our study, we find an increase in Global South authors toward the present, suggesting greater inclusion of local researchers and knowledge-holders. Yet, the rise in Global South authors coincides with the devaluation of taxonomic work by the Global North [77,79,81,85,86]. This dynamic mirrors trends in other professional sectors, in which the devaluation of a profession by hegemonic communities is linked to its labor force being less male and/or less white [87,88], even when the labor (whether physical or intellectual) is regarded as essential. In other words, as a profession becomes more accessible to communities that have been previously marginalized or excluded, the work becomes viewed as less prestigious and is deprioritized in institutional budgets, or vice versa. Given the devaluation of taxonomy as Global North institutions redefine biodiversity research priorities, our findings raise the question as to whether the observed shifts in taxonomic labor reflect a new way in which the Global North exploits essential labor in the Global South.

As sociologist and feminist scholar Joan Acker discusses, labor relations within institutions mirror broader societal dynamics, and institutional practices play a critical role in establishing and maintaining social inequity [89]. In the context of scientific work, and taxonomic work more specifically, explicit and implicit policies dictate who can participate, what roles they take (e.g. finding birds, writing manuscripts, applying for and managing funding, etc.), and how credit is distributed (e.g. first author, senior author, thanked in the acknowledgements, etc.). In many cases these practices differentiate participation in scientific labor along axes of identity, as our results show for nationality, and may also extend to identities like gender, race, and their intersections. These links require us to be more intentional about how scientific work is structured, the roles of different individuals, and how different roles are supported and valued (intellectually and materially).

### (c) The consequences of upholding imperialist structures of power and authority

The observed disparities in eponyms and authorship raise ethical and practical questions about how science is done in a global context. For example, what does it mean for power and authority over the natural world to be disproportionately claimed by the Global North [90,91]? What are the consequences for a community’s relationship to the environment when scientific authority over that environment is held/claimed by individuals outside of the local community [63]? What does it mean for work in the Global South to be translated and consolidated into authority, prestige, and careers in the Global North? Our intention is not to prescribe particular answers to the above questions because formulating those answers will require dialogue between the hegemonic communities which we are part of and those excluded and marginalized by them. Rather, our goal for this paper is to promote conversations and actions around these questions, and contribute to work already being done within and outside the academy that re-frames approaches to science to confront inequity in present-day practices (for examples see [92], https://birdnamesforbirds.wordpress.com, and https://decolonize-dna.org), while acknowledging our collective agency in these practices [93]. As an entry point for these conversations, we recommend working through the questions in the Research Justice Worksheet [94]. We have found this resource helpful for personal reflection and group discussion (thanks, Supriya).

Linnaean taxonomy reflects a social history and practice that continues to consolidate authority in the Global North under the assumption of scientific objectivity. As we grapple with the questions above as a global community, Western science must give up the fallacy of presenting itself as *neutral* and *objective* [95–102], which remain dominant tenets of training and discourse to this day. As sociologist William Jamal Richardson reminds us, “[we] can’t understand the production of knowledge and science independent of its relationship to societal interests and structures of power” [9]. Adhering to the fallacy of neutrality (which is in fact a non-neutral stance and one embedded in white supremacy; [48,102]) has allowed scientists to ignore the social impacts of our actions past and present, while upholding global and institutional structures of dominance and inequity, regardless of intent.

## 5. CONCLUSION

As our findings highlight, access to the Global South continues to be translated into scientific authority, power, and material wealth (e.g. in the form of careers) in the Global North. While we can understand this dynamic by examining the global structures of access and power put in place during centuries of European and U.S. imperialism, a historical perspective alone ignores the agency of institutions and scientists in present-day actions. By ignoring the inequities embedded in Western science, we re-enact and uphold structures of domination and imperialism in our research practices.

## LAND ACKNOWLEDGEMENT

The University of Michigan and the University of Chicago/Field Museum, are situated on the lands of the Council of the Three Fires—comprised of the Ojibwe, Odawa, and Potawatomi Nations—as well as the Miami, Ho-Chunk, Menominee, Sac, Fox, Kickapoo, and Illinois Nations. Our presence as settlers on these lands and our affiliations with these academic institutions have enabled us to write this paper. These institutions—and the US nation that we are a part of—are the products of coercion, theft, ongoing occupation of Indigenous land, and violation of treaties and agreements made with Indigenous groups. For these reasons, we are complicit in the ongoing oppression of peoples indigenous to and displaced from these territories. Along with the seizure of lands, our institutions and practices have driven the erasure and oppression of Indigenous sovereignty and knowledge. We call for our scientific community to: (1) acknowledge and respect Indigenous autonomy and self-determination, (2) reflect on the objectives and impacts of our science, and (3) recognize that our objectives are not justification for the violation of Indigenous autonomy and self-determination.

## ACKNOWLEDGEMENTS

We thank Alvita Akiboh, Alejandra Anchante, Elizabeth P. Anderson, Giorgia Auteri, John Bates, Sasha Bishop, Carlos Daniel Cadena, Susanna Campbell, Alexandra Cook, Harold Eyster, Claudio Gómez-Gonzáles, Eric Gulson-Castillo, Constanza de la Fuente Castro, Michael Lyons, Bruce O’Brien, Diana Macias, Teresa Pegan, Thomas Stewart, K. Supriya, Armando Valdés-Velásquez, Kristen Wacker, Z Yan Wang, Brian Weeks, Ben Winger, Christopher Witt, and Marketa Zimova for their discussions and feedback on the manuscript.

## LINK TO SUPPLEMENT

Versión del manuscrito en español (Spanish language version of manuscript) https://docs.google.com/document/d/1MzxyGtOMW8sYeY7EWg0jKrtxQOzvTDZu/edit?usp=sharing&ouid=116104936007568520737&rtpof=true&sd=true

**Table 1.**
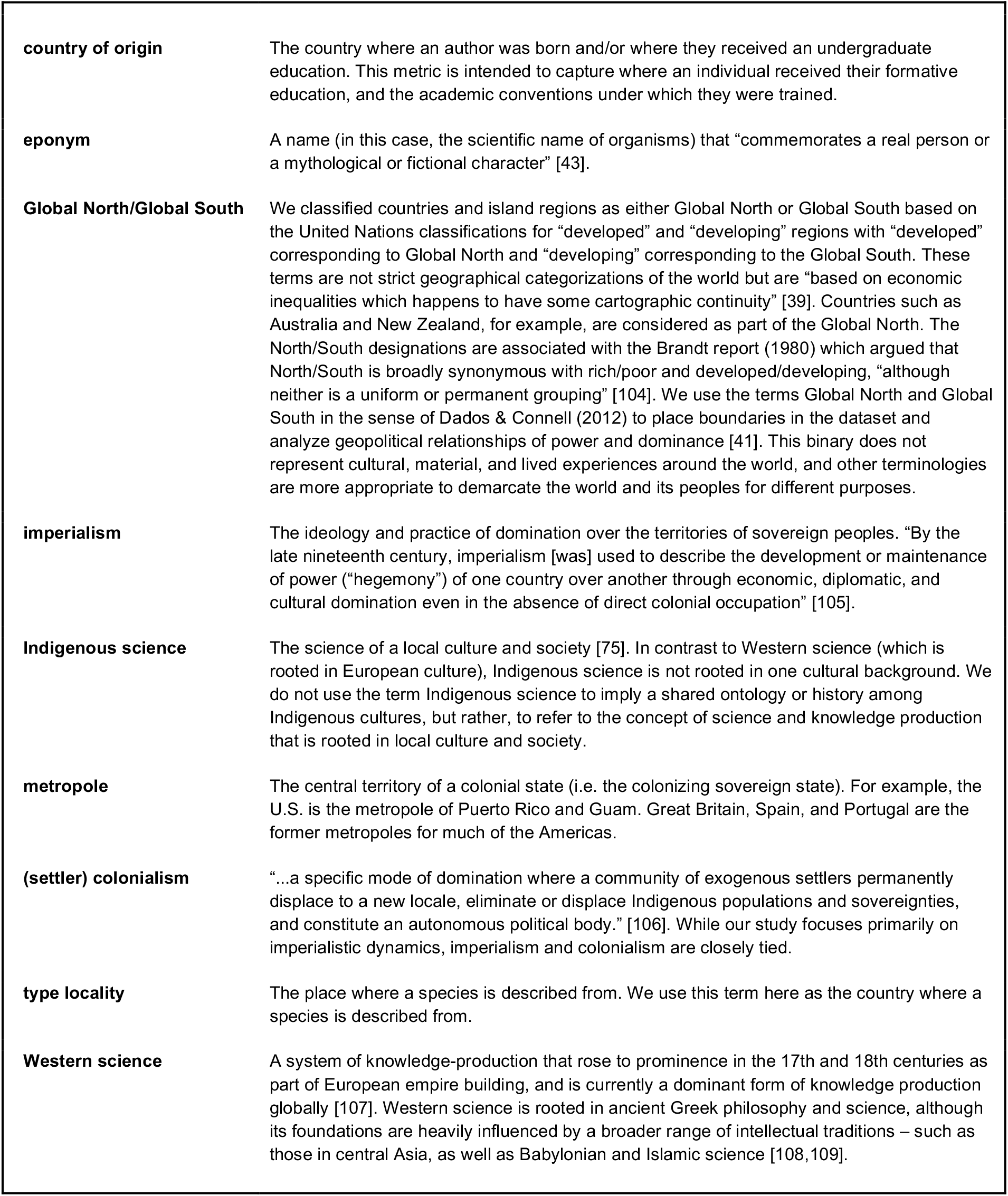
Definitions of terms, and how we use terms in the text.

